# Multi-scale Computational Modeling of Tubulin-Tubulin Lateral Interaction

**DOI:** 10.1101/624213

**Authors:** M. Hemmat, B.T. Castle, J.N. Sachs, D.J. Odde

## Abstract

Microtubules are multi-stranded polymers in eukaryotic cells that support key cellular functions such as chromosome segregation, motor-based cargo transport, and maintenance of cell polarity. Microtubules self-assemble via “dynamic instability,” where the dynamic plus ends switch stochastically between alternating phases of polymerization and depolymerization. A key question in the field is what are the atomistic origins of this switching, i.e. what is different between the GTP- and GDP-tubulin states that enables microtubule growth and shortening, respectively? More generally, a major challenge in biology is how to connect theoretical frameworks across length-time scales, from atoms to cellular behavior. In this study, we describe a multi-scale model by linking atomistic molecular dynamics (MD), molecular Brownian dynamics (BD), and cellular-level thermo-kinetic (TK) modeling of microtubules. Here we investigated the underlying interaction energy landscape when tubulin dimers associate laterally by performing all-atom molecular dynamics simulations. We found that the lateral free energy is not significantly different among three nucleotide states of tubulin, GTP, GDP, and GMPCPP, and is estimated to be ≅−11 k_B_T. Furthermore, using MD potential energy in our BD simulations of tubulin dimers in solution confirms that the lateral bond is weak on its own with a mean lifetime of ~0.1 μs, implying that the longitudinal bond is required for microtubule assembly. We conclude that nucleotide-dependent lateral bond strength is not the key mediator microtubule dynamic instability, implying that GTP acts elsewhere to exert its stabilizing influence on microtubule polymer. Furthermore the estimated bond strength is well-aligned with earlier estimates based on thermokinetic (TK) modeling and light microscopy measurements (VanBuren et al., PNAS, 2002). Thus, we have computationally connected atomistic level structural information, obtained by cryo-electron microscopy, to cellular scale microtubule assembly dynamics using a combination of MD, BD, and TK models to bridge from Ångstroms to micrometers and from femtoseconds to minutes.

## Introduction

Microtubules are dynamic filaments that facilitate critical cellular functions such as chromosome segregation, intracellular cargo transport, and cell architecture. These filaments are composed of tubulin heterodimers, i.e. tightly-associated *α*- and *β*-subunits, with a non-exchangeable site for guanosine triphosphate (GTP) nucleotide binding in the *α*-subunit and an exchangeable site in the *β*-subunit where GTP can hydrolyze to guanosine diphosphate (GDP), followed by release of inorganic phosphate (Pi). This nucleotide exchange at the heterodimer-level confers unique dynamic properties to microtubules, i.e. the stochastic polymerizing and depolymerizing cycles characteristic of dynamic instability (1, 2). The key feature underlying the GTP-tubulin’s greater stability compared to GDP-tubulin, causing alternating growth and shortening phases in microtubule assembly, is yet to be fully understood (3–7). In addition, microtubule dynamic instability is controlled by several factors such as microtubule associated proteins (MAPs) (8, 9), microtubule targeting agents (MTAs) (10–12), microtubule isotype distribution (13), and tubulin post translational modifications (14, 15). These interactions enable microtubules to support important cellular functions (16).

Structural studies of microtubules and tubulin in solution have shed light on conformational states of tubulin heterodimer and how that can explain the different behavior of GTP-vs. GDP-tubulin (17). Those studies revealed that GDP- and GTP-tubulin structures are curved in solution (18–21), compared to a straight structure found in microtubule protofilaments (22, 23), and also, GDP- and GTP-microtubule structure differ by lattice compaction and twist (24). This raises the question of whether tubulin nucleotide state dictates the tubulin preference in curvature predominantly and therefore, their binding efficiency to the microtubule lattice (25). In addition, with recent advances of cryo-electron microscopy (cryo-EM), high resolution structures of tubulin bound to various MAPs and MTAs are now available, revealing different drug binding sites on tubulin (26–28). However, relying only on structural information to explain the regulation of the assembly dynamics by those agents in the context of physiologically relevant problems cannot be achieved. Moreover, acquiring high-resolution structures of dynamic proteins is usually accomplished with using a stabilizing factor (18, 19, 29, 30), which can by itself cause conformational changes to the native structure. Hence, by performing molecular dynamics (MD) simulations of tubulin structures, we are able to study the dynamic evolution of this protein in solution and sample the thermal fluctuations through time. For example, MD simulations of curved structures of tubulin have been used to confirm that they preserve their curvature in solution, with no significant difference at intra-vs inter-dimer bending angles (31, 32). However, using the right time scale and sampling of the ensemble is an important factor in drawing conclusions from the simulation results. In one MD study, using a limited sampling of less than 1ns, the free energy of tubulin for intradimer bending angle was reported, concluding that αβ-tubulin dimers exists in an intermediate bent conformation (33). However, this study did not demonstrate that such a short sampling would represent converged data of the whole ensemble in the equilibrium state. In another study, pushing the limits of MD simulations’s time scale by using a time step of 4fs at the expense of constraining all bonds’ length fluctuations, 3µ simulations of both straight and bent tubulin dimers moved toward a more bent configuration, with GTP-tubulin showing a wider range of bending flexibility compared to GDP-tubulin (34).

Considering the limits of current computational resources, capturing kinetic information from MD simulations at a time scale of ~*μs* remains prohibitive, especially considering that multiple replicates are needed to sample the distribution of initial conditions. By contrast, coarse-grained Brownian dynamics (BD) with ~ms time scales at a cost of atomistic detail, and thermo-kinetic (TK) simulations with less detail allowing access to ~100s time scales, together enable recapitulation of the kinetic rates and microtubule tip structures consistent with those found *in vitro* and *in vivo* (35–38). The interactions of particles in the BD simulations, here being the tubulin dimers, are modeled using an input potential, which can be dissimilar for tubulin as a function of its nucleotide state in different studies (7, 38–41). However, the interaction energy profiles have been adjusted in these models to match experimental observations of MT tip structure, dynamic assembly rates, and microtubule stiffness. Therefore, this discrepancy in potential of interactions in different BD models has resulted in incompatible conclusions about various aspects of microtubules’ behavior. For instance, catastrophe, microtubule’s switching from growth to shortening, has historically been described as stemming from loss of the stabilizing GTP-cap due to hydrolysis (37, 42–44), while recent Brownian dynamics studies (39, 45) suggested that stochastic variations in the number of curled protofilaments is responsible for causing catastrophe, albeit while using a different shape and strength for potential of interactions of tubulins from previous models (41, 46). In addition, lateral bond formation has been identified as the limiting step in forming the microtubule lattice with a large entropic component in another study (40), but no direct calculations have been reported for the lateral bond strength or its nucleotide-state dependency even though it is critical for assembly and potentially for dynamic instability as well.

In this study, the questions that we address are: 1) what role does the lateral bond play in MT assembly and stability, and 2) more generally, how do we connect modeling simulations from atoms up to organelles, such as microtubules, at the cellular level. We addressed these questions by developing a multi-scale approach to study tubulin-tubulin interaction, building a framework for more complex interactions with MTAs and MAPs. We use high resolution cryo-EM structures of tubulin for initiating full-atom molecular dynamics simulations to study the dynamic evolution of different nucleotide states of tubulin in solution. Using the equilibrated structure, we then calculate an energy landscape for tubulin-tubulin lateral interaction in terms of a potential of mean force (PMF), using multiple replicates. To our knowledge, such free energy calculations on a large globular protein-protein system (MW=110 kDa per heterodimer), ~200,000 atoms, using full-atom MD simulations has not been reported. This enabled us to probe for possible energetic differences between GTP- and GDP-tubulin in assembly dynamics, and whether it is the lateral bond strength that distinguishes the two states in terms of establishing microtubule stability. Furthermore, we used the PMF obtained as an output from molecular dynamics to define the input potential energy in Brownian dynamics simulations, as previously developed by Castle et al. (38). Overall our work provides a multi-scale modeling approach (Figure 1) to use interaction energy profiles from full atom MD simulations in Brownian dynamics, which we have previously linked to a thermo-kinetic model (37), and so we are now able to establish a framework for moving from crystallographic/cryo-EM structures to MT assembly behavior. Assembly dynamics can be used furthermore in cell-level (CL) modeling of the microtubules (47) to predict more complex physiologically-relevant behavior such as the spatiotemporal distribution of MTs within the cell. Such a multiscale MD-BD-TK-CL framework is able to seamlessly connect protein dynamics at femtosecond time scales and Ångstrom length scales to entire cell level behavior with an ensemble of 100’s of microtubules at minutes-hours time scales and micrometer length scales (36–38).

**Figure 1.**
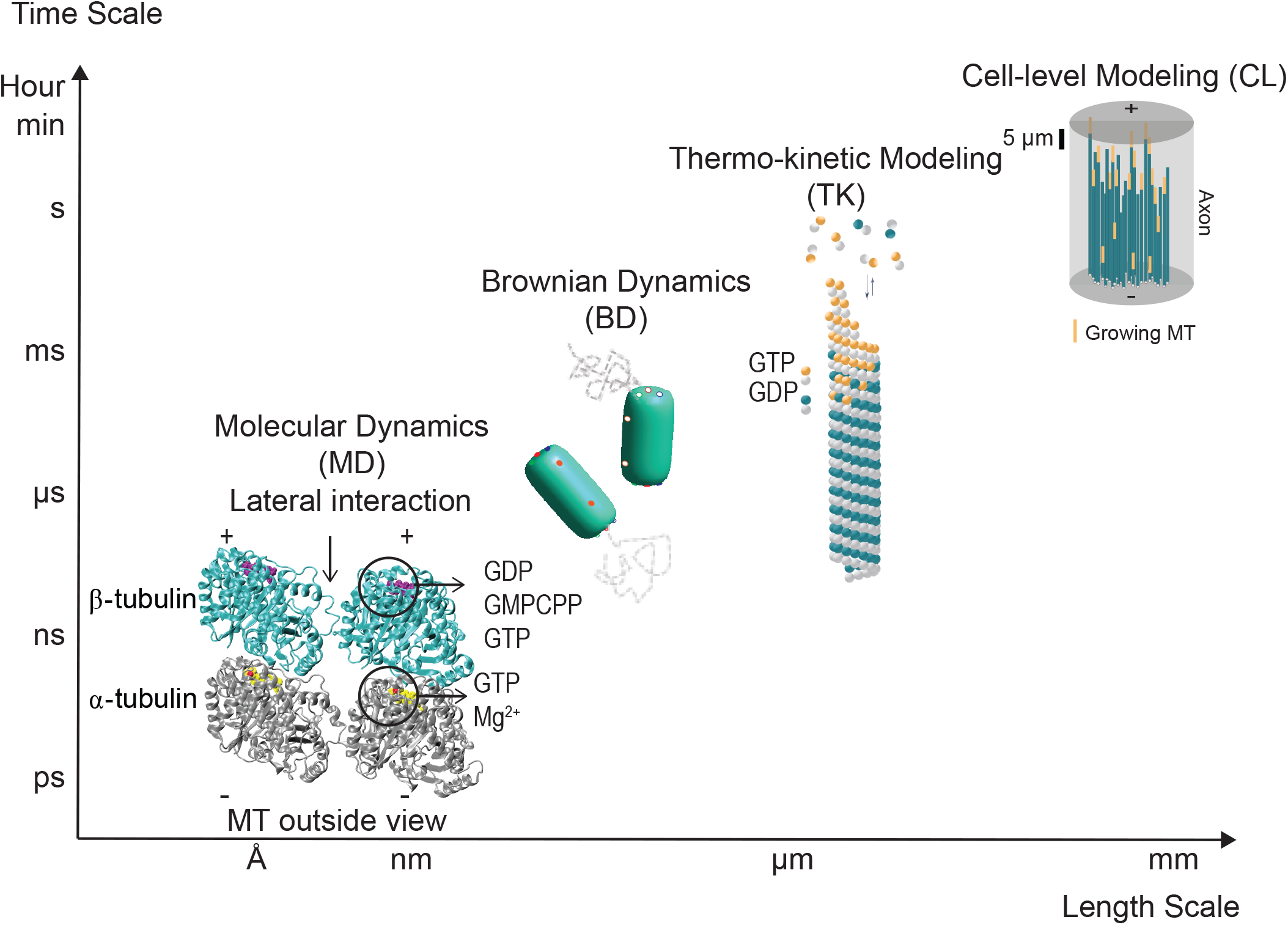
Illustration of the multi-scale approach to studying microtubule self-assembly at various length-time scales, including molecular dynamics (MD) (PDB ID: 3JAS, 3JAT), Brownian dynamics (BD), thermo-kinetic (TK) modeling and cell-level (CL) modeling.

## Methods

### Simulation system preparation

Our computational simulations focused on tubulin heterodimers with one lateral neighbor in three different nucleotide states, i.e. GDP, GMPCPP, and GTP. However, tubulin in the GTP-state is unstable due to its tendency to hydrolyze with a rate constant of ~0.1-1.5 s^−1^ (36, 46, 48). This makes it difficult to obtain a crystal structure of this state without an additional stabilizing element such as RB3-stathmin-like domain and DARPIN protein (18, 19). For our study, the three dimensional GDP- and GMPCPP-state tubulin structures with one lateral neighbor were extracted from the published cryo-EM dynamic structures of microtubule by Zhang et al. (29) (PDB ID: 3JAS, 3JAT). The structures were obtained in the presence of kinesin head domains decorating the microtubule lattice to distinguish between α- and β-tubulin subunits while presumably having little effect on the microtubule structure. We then built our GTP-state system modifying the initial structure of the GMPCPP-state tubulins and equilibrating the total complex. In each state, we have two tubulin dimers laterally paired, as they would in a microtubule lattice, with the GTP-associated Mg^2+^ present. The systems were then separately solvated in TIP3P water (49) using an 8Å clearance from each side, resulting in a periodic cubic box with dimensions of 125 × 90 × 124 Å, on average. MgCl_2_ ions were added at 2mM concentration to neutralize the system (31 Mg^2+^ and 2 Cl^−^) based on physiologically-relevant salt concentrations. A total number of 128,500 to130,000 atoms was used in all the systems.

### Molecular Dynamics Simulations

MD Simulations of all three nucleotide systems were run using NAMD 2.10 software (50) using the CHARMM 36 force field (51). The protein complex along with the nucleotides were all parametrized using the CHARMM-GUI interface (52). Each simulation system was initially energy minimized for 12000 steps using the conjugate gradient algorithm. The systems were then solvated and neutralized with MgCl_2_ at 2mM. The solvated systems were heated to 310 K for 1ns using a Langevin thermostat (53), and then run for 7ns in an NPT ensemble (T=310K and P=1 atm) with the backbone atoms of the proteins being initially constrained with a harmonic potential in all directions having a spring constant of k=2 kcal/mol/Å^2^, and then gradually decreasing the harmonic constraints by dividing the spring constant in half in each 1ns run to prevent large fluctuations from occurring. The simulations were followed by a total production run of 150ns for each system (equilibrating runs). All simulations were run with 2 fs time step and a cutoff radius of 12Å for van der Waals interactions, using Particle Mesh Ewald (PME) for long range non-bonded interactions (54). NVIDIA Tesla K40 GPUs were used to accelerate the simulations on the Mesabi cluster at the Minnesota Supercomputing Institute (MSI), University of Minnesota, and NVIDIA Kepler K80 GPUs were used on Comet, an Extreme Science and Engineering Discovery Environment (XSEDE) (55) dedicated cluster at the San Diego Supercomputing Center (SDSC).

A second set of simulations were run for free energy calculations. Running our system for ~200ns did not yield a full energy landscape of tubulin-tubulin interactions due to insufficient sampling of progressively less bonded (higher energy) states and existence of possible local minima in the energy profile. Since overcoming those energy barriers is beyond that afforded by MD simulations, we employed the umbrella sampling method (56) to sample the ensemble sufficiently and have independent simulations that each can be run for longer sampling time in parallel, considering the large number of atoms. Consequently, this method yields a better convergence compared to the adaptive biasing forces (ABF) method (57). We obtained a potential of mean force (PMF), a free energy landscape as a function of a specified reaction coordinate. For investigating the lateral potential of interaction, we defined the lateral center-of-mass to center-of-mass distance of the dimers to be the reaction coordinate, as further verified to be the most probable path of unbinding for two similar tubulin dimers as obtained from Brownian Dynamics (see Fig. S1 in the Supporting Material). The bias potential stiffness was tuned to be 10 kcalmol^−1^Å^−2^ to give sufficient overlap of the histograms of the windows while not being too soft resulting in a large correlation of dimers’ movements (see Fig. S2 in the Supporting Material). To cover the full range of inter-dimer interactions, 18 windows were created, each being 1Å separated from their nearest window. The production run was used to choose 10 equilibrated initial structures for creating replicates of umbrella windows for each system. The structures were selected far enough apart at time points after where the backbone RMSD plateaued in the equilibrium run (~50ns) and separated by the correlation time of the dimers’ movement (10-20ns for different simulations). Having multiple replicates reassured us that the resultant time averaged PMFs are converged to the ensemble-averaged interactions. Each window was then equilibrated for 10ns constrained by the bias potential and followed by a 20ns sampling run for free energy calculation. For determining the convergence of the PMF for each replicate, we increased the sampling time incrementally (by 5ns), calculated the PMF, and compared it to the previously calculated PMF to ensure that the change of energy does not exceed a threshold of 1.5 k_B_T, determined by the Monte Carlo bootstrap error analysis of the PMFs (see Fig. S3, Table S1 in the Supporting Material). To evaluate the effect of salt concentration on the lateral interaction of tubulin, a third set of simulations were run with GDP-tubulin in a neutralized system with 2mM MgCl_2_ and an additional 100nM of KCl. Three replicates of PMFs were calculated and the average was compared to the PMF obtained from the initial neutral system obtained from ten replicates (see Fig. S4 in the Supporting Material).

### Analysis of simulation trajectories

The equilibrium run trajectories were stored every 3000 time steps (6 ps) and the reaction coordinate in free energy simulations was recorded every 200 time steps (0.4 ps). The stored trajectory files were analyzed for several conformational changes, such as RMSD and a detailed interaction energy decomposition. The software VMD 1.9 was used for visualization of the trajectories (58). Weighted histogram analysis method (WHAM) (59, 60) was used to combine the histograms and build the unbiased PMF in a memory-efficient way.

For analyzing the equilibrium trajectories, the buried SASA can be calculated as summing over the SASA of the two dimers separately minus the SASA of the laterally-paired tubulins, divided by two, given as

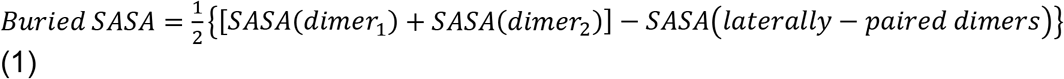

The hydrogen bonds and salt bridges were calculated with the plugins available in VMD.

### Brownian Dynamics Simulations

We examined the kinetics of tubulin dimers’ lateral interaction according to the Brownian dynamics model of Castle *et al.* (38) with the modification of simulating two dimers in solution as opposed to a single dimer binding to the microtubule lattice. In the BD simulations, only lateral association/dissociation is possible and the potential energies are the entropy corrected PMFs from MD simulations. Simulations were run for three nucleotide cases and each was run for 500,000 iterations of binding simulations (total time varying from 0.1 μs −1 ms) and 50,000 iterations of unbinding simulations (total time 0.5 ns – 5 μs). For binding runs, a distance criterion was defined based on the input potential of interaction, where two subunits are considered “bound” laterally if all of the lateral zones’ distances are within the binding radius (*r*_*i*_ ≤ *r*_*b*_). Binding radius for PMF energy profiles was defined where the slope (inter-particle force) reached 50% of its maximum value. Considering the similarity of the PMFs to a Lennard-Jones (LJ) potential, the force was close to maximum near the minimum of the potential and was very low where the potential plateaus to zero. Therefore, this definition of r_b_ ensured us that the bound particles felt a force significant enough to hold them together with a high probability (90%). For unbinding simulations, we used a separation distance criterion of R_U_=11nm, according to (38), where the probability of rebinding is very low (p <0.01). Additionally, we ran BD simulations of perfectly aligned tubulin pairs starting from the minimum of the potential interaction energy and investigated the unbinding path, extent of tubulin rotation upon unbinding, and dissociation time to compare to MD simulations’ reaction coordinate and time scale.

## Results and Discussion

### Equilibration of tubulin dimers in varying nucleotide states predicts a lack of nucleotide sensitivity of dimer structure in solution

To investigate the nature and strength of the lateral interaction between two tubulin dimers using our multi-scale approach, we first built a molecular dynamics model of two tubulin dimers laterally paired with one of three nucleotide-states: GDP-, GMPCPP-, or GTP-tubulin. Similar to earlier molecular dynamic studies of tubulin heterodimers (32, 33, 61), our simulations (~200ns vs. 10-100ns equilibration in previous studies) indicated that αβ-tubulin structure is stable and consistent in aqueous environment regardless of the nucleotide state or lateral neighbor (see Movie S1-S3 in the Supporting Material). Mean root-mean-squared displacement (RMSD) of backbone atoms was measured from the starting coordinates obtained from the cryo-EM structure as the protein structure stability for three nucleotide states of the dimer. We equilibrated both a single tubulin dimer in solution and laterally-paired tubulins for 200ns (Figure 2A, B). The asymptotic (plateau) behavior of the backbone atoms’ RMSD after ~50ns for all conditions without any significant fluctuations reveals that the initial cryo-EM structures are not far from the equilibrium states for single dimers or pairs of laterally-associated dimers in solution. In addition, having created the GTP-tubulin structure from GMPCPP-tubulin structure, the RMSD trend confirmed that the estimated GTP-structure was stable. To identify the conformational differences between the backbone atoms of different nucleotides, we performed a residue-to-residue RMSD comparison for the average structure using the 150ns of the production run of different nucleotide-states of tubulin (Figure 2C). It was observed that the residues in flexible loops have the highest RMSD in each case and the average position of main helices and beta sheets of the backbone atoms remain relatively unchanged among different nucleotide states. Additionally, we examined the flexibility of the residues for the laterally-paired tubulins and observed that the M-loop involved in the lateral bond has significantly lower RMSF compared to the free M-loop on the neighbor dimer and that the M-loop in α-tubulin is more ordered compared to β-tubulin. Hence, the only major conformational difference between the laterally-paired and a single free dimer was demonstrated to be the ordering of the flexible loops at the lateral interface rather than the backbone structure. Thus, we conclude that the backbone structure of the tubulin heterodimer as measured from cryo-EM measurements is very similar to that predicted from MD at equilibrium for a single dimer or pairs of laterally-associated dimers in solution, independent of nucleotide state. This further provides justification for investigating lateral and longitudinal interactions of tubulin by simulating only two tubulin dimers in solution without having to simulate the whole MT lattice.

**Figure 2.**
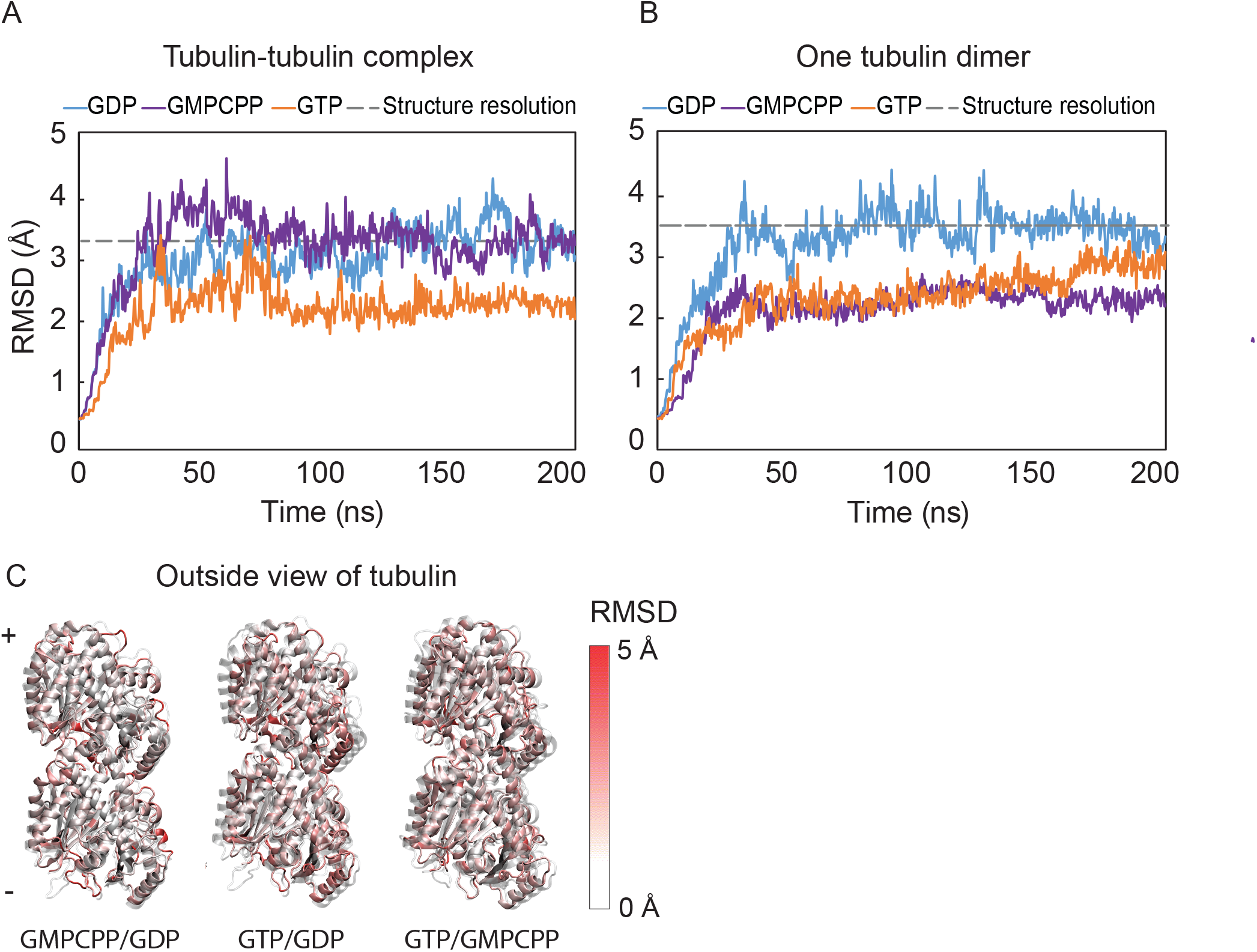
Stability of the backbone atom structure when equilibrated with water and ions is shown as the average root-mean-squared deviation (RMSD) of the protein system, for tubulin-tubulin complex (A) and tubulin dimer backbone atoms (B) during the equilibration with water and ions. RMSD per residue for the comparison of different nucleotide’s average structure is shown in (C) for GMPCPP compared to GDP, GTP to GDP, and GTP to GMPCPP respectively.

### Interaction energy decomposition at the tubulin-tubulin lateral interface indicates that lateral interactions are largely independent of dimer nucleotide state

To analyze the contribution of different types of non-covalent interactions predicted to mediate the lateral bond between adjacent tubulins, we decomposed the lateral inter-dimer interaction energy into H-bonds, ionic, and hydrophobic interactions. The equilibrium simulations of each nucleotide state were analyzed for each of these three types of interaction, accounting for the autocorrelation time between each data point of the trajectories to ensure that we obtained temporally-independent samples (Figure 3). On average, ~6 inter-dimer H-bonds were found among all three cases (Figure 3A). The number of inter-dimer salt bridges which were present for more than half of the time trajectory, were calculated to be ~2 on average, indicating an important contribution of ionic interaction between two lateral neighbors (Figure 3B). GMPCPP- and GTP-tubulin showed slightly stronger ionic interaction at the dimer interface compared to GDP-tubulin. For the last component, hydrophobic interactions, we calculated the inter-dimer buried solvent accessible surface area (SASA) as the hydrophobic pocket that would have weak interactions with water. Figure 3C shows that unlike the ionic interactions, GDP-tubulin has a slightly higher inter-dimer hydrophobic interaction compared to the other two states. Important residues involved more than 25% in lateral interaction are shown in Table 1. There are two main favorable interactions found within the lateral interface indicated in Figure 3D (see Movie S1-S3). A lock-and-key interaction between M-loop (the key) on one side and H2-S3 and H1’-S2 loops (the lock) on the other side in both subunits, which was also identified as the main lateral contacts in the cryo-EM study (29), and an interaction between H3 on one side and H9 and H9-S8 loop on the other side, which was not found within the cryo-EM structure, are identified from our results. Our analysis indicates that the interaction between H3 and H9-S8 loop is the strongest in all nucleotide cases and the interaction between M-loop and H1’-S2 is the weakest (<25% in all states). This finding highlights the importance of studying the interactions over the course of a dynamic simulation where side chains are allowed to thermally fluctuate and interact with the neighbors in contrast to a vitrified protein structure used to obtain cryo-EM structures. Since the M-loop is relatively ordered in the laterally bonded state, compared to a single dimer in solution, we conclude that the decreased entropy upon binding acts to destabilize the lateral bond (40). Besides, the decreased fluctutations due to lateral bond is more amplified in the M-loop in β-tubulin, highlighting a key lateral role for the M-loop in β-tubulin compared to α-tubulin (62). In addition, through the inter-dimer interaction decomposition analysis that different nucleotide states have only modestly dissimilar contributing energy components in stabilizing their lateral bond. Even so, the total strength of the interaction components taken together as lateral bond remains to be calculated and cannot be determined with confidence through equilibrium simulations only.

**Table 1.** Important interface residues identified in lateral interaction of the tubulin dimers

| Inter-dimer interaction | Important involved residues (>25% occupancy) | Secondary Structure | Tubulin state (>25% occupancy) |
| --- | --- | --- | --- |
| H-bonds | LYS124-ASP297 ($\beta$) | H3--H9 | all |
| | LYS299-ASP120 ($\beta$) | H3 -- H9-S8 | all |
| | SER128-GLU290 ($\beta$) | H3 -- H9-S8 | GDP (40%) |
| | ARG308-ASP116 ($\beta$) | H3 -- H9-S8 | GMPCPP (26%) |
| | ARG308-ASP127 ($\alpha$) | H3 -- H9-S8 | GMPCPP (30%) |
| | TYR283-GLN85 ($\beta$) | M-loop -- H2-S3 | GDP (27%), GMPCPP (26%) |
| | ARG123-GLU297 ($\alpha$) | H3 -- H9-S8 | GMPCPP (26%), GTP (25%) |
| Salt bridges | ASP120-LYS299 ($\beta$) | H3 -- H9-S8 | all |
| | GLU127-LYS338 ($\beta$) | H3 -- H10-S9 | all |
| | ASP127-LYS338 ($\beta$) | H3 -- H10-S9 | GMPCPP (93%), GTP (31%) |
| | GLU90-LYS280 ($\alpha$) | M-loop -- H2-S3 | GDP (31%), GTP (28%) |
| | GLU290-LYS124 ($\beta$) | H3 -- H9 | GMPCPP (40%) |
| | ASP116-LYS299 ($\alpha$) | H3 -- H9-S8 | GMPCPP (30%) |
| | GLU284-LYS124 ($\alpha$) | M-loop -- H3 | GMPCPP (26%) |
| | GLU297-LYS124 ($\alpha$) | H3 -- H10-S9 | GMPCPP (35%) |

**Figure 3.**
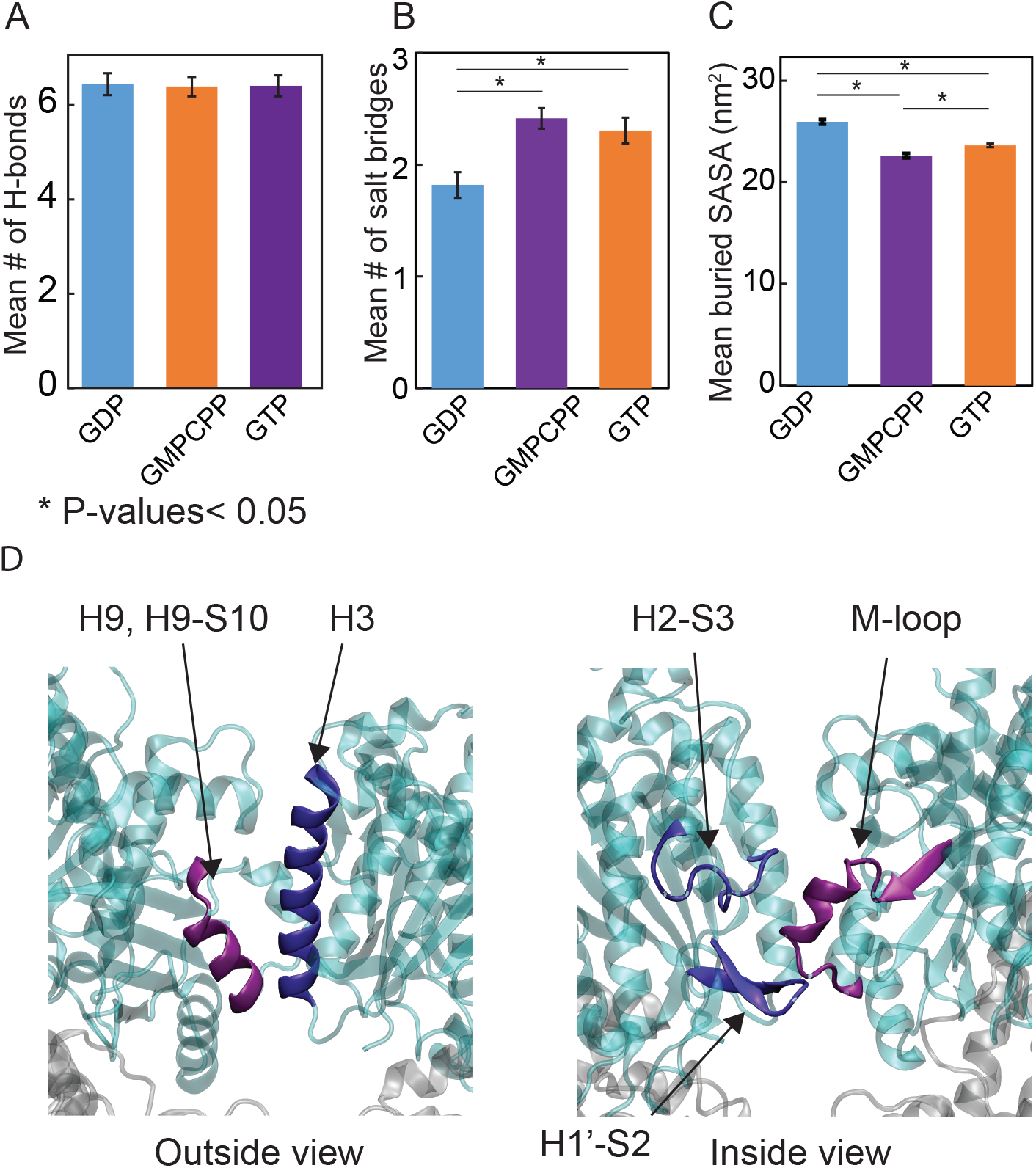
Inter-dimer lateral energy decomposition does not depend on nucleotide state over the production run (150ns), including mean number of inter-dimer hydrogen bonds (A), mean number of inter-dimer salt-bridges (B), and mean solvent-accessible surface area (SASA) (C). The main interface residues involved in the lateral interaction are shown in (D). Error bars indicate standard error of the mean (SEM) of 100 bootstrap sets of data considering the correlation time of the time series.

### Tubulin dimers exhibit only modest intradimer bending as a function of nucleotide state

Since our starting structure for our simulations was obtained from straight protofilaments through cryo-EM (29), we were interested to see whether the structure of the tubulin dimers deviated from the lattice structure once they were equilibrated in solution without any constraints. In particular, microtubule stability has been proposed to depend on the straight conformation of tubulin dimers, while instability has been proposed to result from bent tubulin. This led us to calculate the conformational difference of the three nucleotide states in terms of the intra-dimer angle between the individual α- and β-subunits (18, 19, 33).

To assess whether nucleotide state and the absence of the microtubule lattice influences intradimer bending angle, we calculated intra-dimer bending angle for equilibrated dimers as the relative angle of the β-subunit to the α-subunit from the last 150ns of the MD production run after equilibrium. The rotation angle was calculated as the angle between the vector connecting the center-of-mass (COM) of α- and β-subunits at time zero (crystal structure) and time t, after superimposing α-subunit backbone atoms (Figure 4A-B), similar to the methodology in previous studies (31, 33, 63). We decomposed the angles of rotation into three perpendicular bending modes (Euler angles) rather than calculating the dot product of the two vectors since we can obtain more information about the directions of bending using this approach. The angle decomposition into x-, y-, and z-directions (Figure 4C), indicates that there is almost no intra-dimer twist (θ_z_), only a small tangential bending (θy ≅1-2º), and the majority of rotation happens around bending radially outward (θ_x_). This result is consistent with previous MD simulation (~15ns) of tubulin dimer bending both tangentially and radially outward in solution, nucleotide-independent (34, 63, 64) (referred to as twist-bend mode in (34)). In addition, bending angles calculated for the laterally-paired heterodimers in solution revealed that the presence of a lateral bond diminshed the intradimer bending of tubulins, independent of nucleotide, in line with the results of Peng *et al.* (2014). Although the structures of both GDP- and GTP-tubulin move slightly toward a bent conformation and are significantly different from the crystal structure, the extent of the outward bending (~3º) does not match the 12-15° intradimer rotation observed when tubulin is bound to stathmin domain and DARPin protein or in γ-tubulin structure obtained from small-angle X-ray scattering (SAXS) (18, 19, 65). We conclude that the discrepancy of bending angle between our simulations and stathmin/DARPin-bound tubulin may stem from the fact that this degree of freedom is slow to relax (high correlation time of ~10-30 ns during 150ns of the production run) and further sampling time is required to reach a conclusion about the final converged mode of bending. This is further confirmed by comparing our data to the initial time data points of modest bending motion in Igaev *et al.* (2018) (figure 3-figure supplement 1), and the data in the coarse-grained simulations of protofilaments in Grafmuller *et al.* (2011), suggesting that further simulation time is required for time convergence of the angle data. Even so, having simulated tubulins for several µs, Igaev *et al*. did not see a bending convergence between the curved and straight conformations of unpolymerized tubulin, indicating that experimentally obtained curved conformations do not fully reveal the bending mode of pure unpolymerized αβ-tubulin.

**Figure 4.**
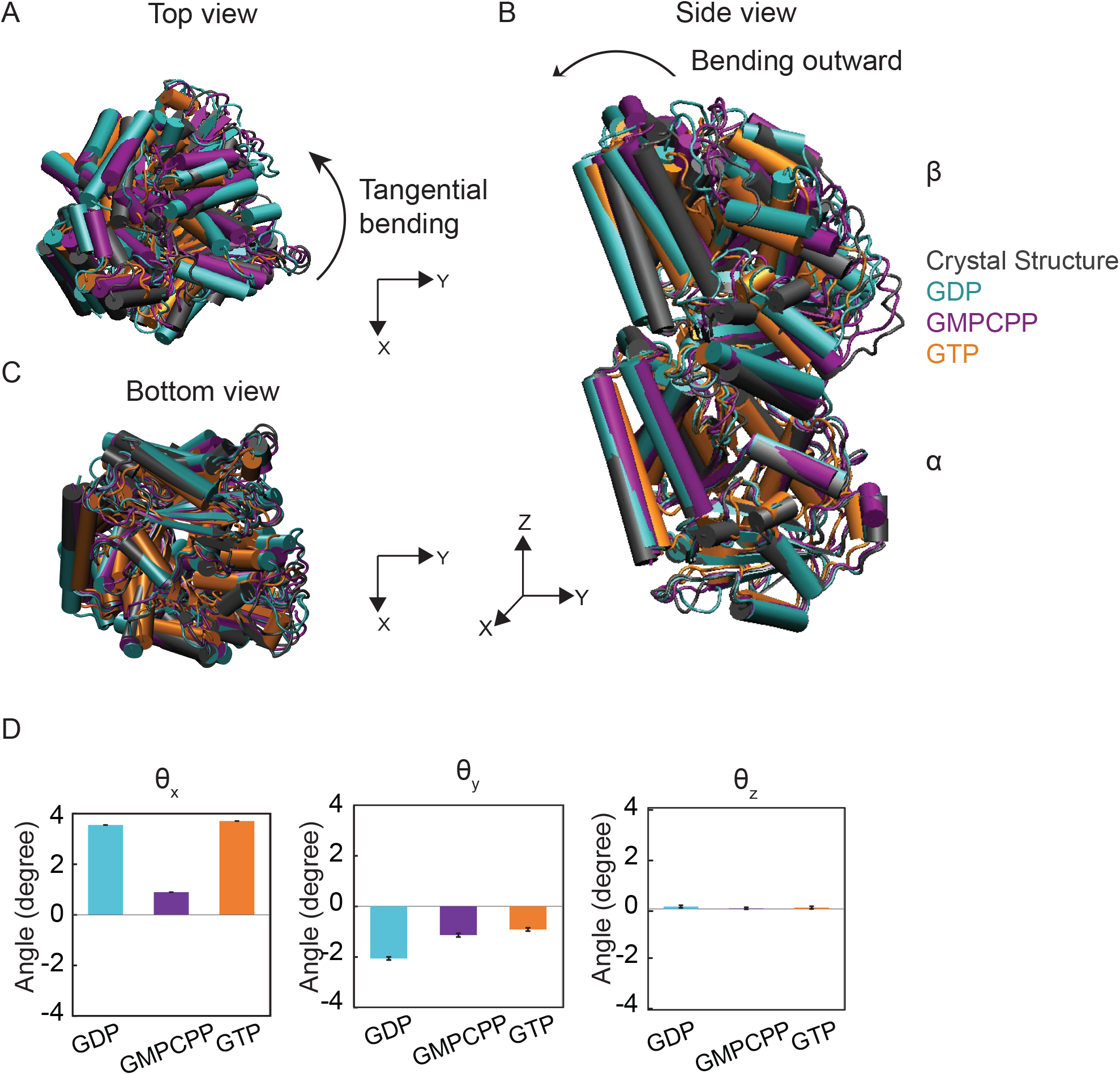
Intradimer angle rotation of tubulin dimer in solution shows both GDP- and GTP-tubulin move toward a slightly bent configuration. Top view of tubulin (A), side view of tubulin (B), bottom view of tubulin (C) and quantitative values of the angles decomposed in x, y, and z axis (D). Cyan, purple and orange indicate GDP-,GMPCPP-, GTP-tubulin average structure, respectively, from MD simulation, and grey is GDP-tubulin crystal structure. Angles are measured as rotational angles required to superimpose the monomer vector connecting center-of-mass of β- and α-H7 helices to its reference crystal structure vector after aligning the whole dimer based on the α-subunit. Error bars indicate standard error of the mean (SEM) of 100 bootstrap sets of data considering the correlation time of the time series.

### Different nucleotide-state tubulins do not have significantly different lateral bond free energy landscapes

Equilibrium MD simulations and simple Boltzmann sampling are not efficient approaches to estimate the total energy landscape as two dimers separate laterally. Therefore, we sought to calculate the total lateral interaction energy by performing free energy calculations to obtain a potential of mean force (PMF) as a function of nucleotide state (Figure 5A). Umbrella sampling (56) was used for our simulations due to its ability to sample parallel windows independently and more flexible sampling time to ensure the convergence of the calculations. We performed the umbrella sampling method for a center of mass-to-center of mass between dimers’ distance reaction coordinate, which we determined is the most efficient binding path according to BD simulations (Fig. S1). We optimized the umbrella sampling method for the best fit biasing potential (Fig. S2), the sampling time, and the PMF convergence along ten distinct replicates (Fig. S3 and Table S1). As shown in Figure 5A, we observe that there is no energy barrier found between the bound and unbound states in any of our PMF replicates and is unlikely to exist because there is no significant desolvation of the binding interface, in contrast to the assumption made in some previous BD studies (39, 40). For comparison, the PMF obtained by MD is plotted along with previously published BD estimates of the lateral bond strength (38, 39) (see Fig. S6 in the Supporting Material). A statistically significant (p<0.005) minimum shift is observed between the GDP-state being at 53.7 Å and the average minimum of GMPCPP- and GTP-states being at 52.4 Å. This modest shift is only important when there is incongruity in the lattice, i.e. an ensemble of GDP- and GTP-tubulins with different preferred lateral distance, which can appear as an existing mechanical strain stored in the lattice (66). In a heterogeneous lateral interface, both GDP- and GTP-tubulin will settle down on an intermediate minimum distance, which will be equal to ~1 k_B_T energy change in the lateral bond strength according to both PMFs. Moreover, we incorporated this minimum shift into our BD simulations of tubulin dimers both in the solution and in the lattice (using the lateral potential from our MD simulations and our previously estimated longitudinal energy profile for the lattice assembly (38)) to investigate if this shift affects the kinetics of the dimers (see Table S2 and Fig. S7 in the Supporting Material). No significant change was detected between the associations of GDP- and GTP-tubulin in solution and the dissociation rate was found to be high (~10^7^ s^−1^) in both cases, with GDP-tubulin having slightly weaker lateral bond energy 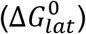. We conclude from the BD results that this minimum shift (~ 1 Å) appears only in the ensemble of GDP- and GTP-tubulin lateral bond and is tolerated within the MT lattice.

**Figure 5.**
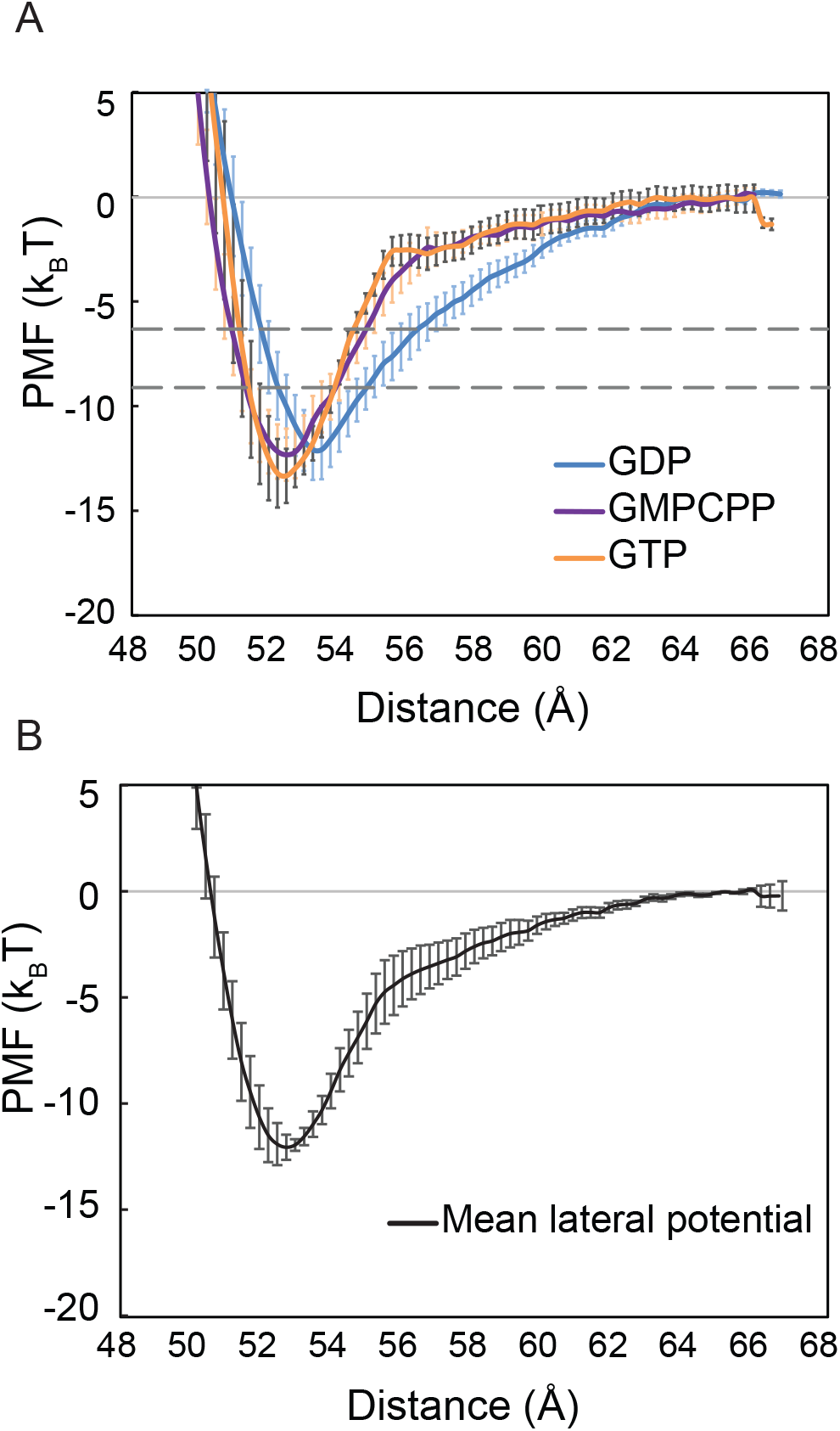
Mean potential of mean force (PMF) of tubulin-tubulin lateral interaction is found to be nucleotide-independent for three nucleotide states, GDP-(blue), GMPCPP-(purple) and GTP-tubulin (orange) (A). Dashed lines are showing the lower and upper estimates of lateral interaction well depth (U_lat_) from previous publications (39, 77). The error bars are the standard error of the mean (SEM) of ten replicates for each nucleotide. Average lateral PMF of the three nucleotide states is shown in (B). The error bars are SEM of the three states.

PMF replicates obtained for the three different nucleotide states were then analyzed for their well-depth, shape (energy’s first spatial derivative during unbinding), and binding radius (Table 2). We defined a half-force radius as an indicator of where the derivative of the energy profile (force) decays to 50% of its maximum value when unbinding occurs (Methods section). The binding radius at which we see significant interaction between the dimers is around 0.5 nm (half-force radius) which is close to the value of Castle *et al*. 2013. Well-depths were also defined as the difference between the average of ten points around the minimum and 10 points around the maximum of the profile since the dimers have thermal fluctuations when bound or unbound. The well-depth values are found to be higher than the harmonic potential strengths previously used in *Castle et al.*, however, considering the shape of our PMFs being closer to a Lennard-Jones potential, with a slow decay rate to zero compared to a steep decay of a harmonic potential, the ultimate kinetic rates are found to be well aligned. Interestingly, by this metric the well depth, binding, and half-force radius of the PMFs were not statistically different from each other as a function of the nucleotide state, and therefore, we are able to calculate one final average PMF for the lateral interaction of tubulin regardless of the nucleotide state using all 30 replicate simulations (Figure 5B). Additionally, to assess the influence of ionic strength, we increased the salt concentration in the solution and ran 3 replicates of GDP-tubulin solvated with an additional 100nM KCl (Fig. S4). The analysis of the PMF profile indicates a slight but statistically significant (p < 0.02) minimum shift from 5.39 nm to 5.19 nm in the KCl added solution. All other properties of the profiles, including the well depth, binding and half-force radius were not statistically significant (Table S3). We conclude that adding more salt in the solution mediates more interactions at the interface, bringing the subunits closer together at the cost of disrupting previous interactions, such that the total lateral bond strength remains the same.

**Table 2.**
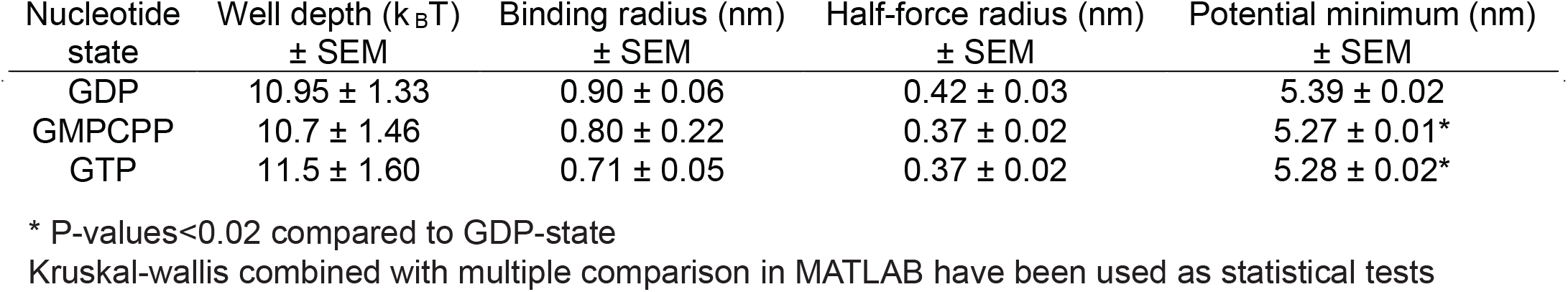
Potential well-depth values with associated SEM (of ten replicates for each nucleotide state), potential minimum location and calculated binding and half-force radii for three simulation cases. Values are calculated based on the average of ten data points around the absolute maximum or minimum due to thermal fluctuations.

| Nucleotide state | Well depth ($k_B T$)<br>± SEM | Binding radius (nm)<br>± SEM | Half-force radius (nm)<br>± SEM | Potential minimum (nm)<br>± SEM |
| --- | --- | --- | --- | --- |
| GDP | $10.95 \pm 1.33$ | $0.90 \pm 0.06$ | $0.42 \pm 0.03$ | $5.39 \pm 0.02$ |
| GMPCPP | $10.7 \pm 1.46$ | $0.80 \pm 0.22$ | $0.37 \pm 0.02$ | $5.27 \pm 0.01^*$ |
| GTP | $11.5 \pm 1.60$ | $0.71 \pm 0.05$ | $0.37 \pm 0.02$ | $5.28 \pm 0.02^*$ |

\* P-values<0.02 compared to GDP-state
Kruskal-wallis combined with multiple comparison in MATLAB have been used as statistical tests

Previous studies modeling MT dynamics have used different potentials of interaction for tubulin (7, 37–39, 41, 46, 67, 68). We reviewed the values found for lateral and longitudinal bond strengths and the energy difference distinguishing GDP- and GTP-state (∆∆*G*°) in Table 3. Note that in Brownian Dynamics simulations, there is a difference between the intrinsic bond energies (U) and the total bond energy 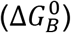, mainly because the interaction zones are not all perfectly aligned when two subunits approach and bind to each other. Therefore, 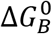 would be the time average of lateral interaction energy and not the maximum intrinsic energy (well-depth), a distinction described previously by Caste et al. 2013 (38). Since MD energy values are not absolute values and have a standard deviation around the mean due to variability in the initial conditions sampled from equilibrated structures, random sampling within the standard error of the mean is required. We sampled through entropy-corrected PMFs within their standard error of the mean as the input for our BD simulations and calculated 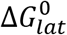 for GDP- and GTP-state and a series of ∆∆*G*° values. We calculated the probability at which a specific ∆∆*G*° used in previous models can be found within the ∆∆*G*° values that resulted from our BD simulations. The analysis indicates that a ∆∆*G*° larger than 2 k_B_T is not consistent (> 95% confidence) with the 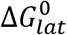 values from the two energy landscapes obtained in MD simulations. Overall, our results imply that the GTP/GDP-state dependent energetics driving the dynamic instability behavior of microtubules are not related to nucleotide-dependence of the lateral interactions. This is consistent with the observation from the cryo-EM structure comparison of GDP- and GMPCPP-protofilaments (29), that the main structural difference between the two nucleotide states resides in the longitudinal compaction of the lattice rather than at dimers’ lateral interface.

**Table 3.** Estimates of tubulin lateral and longitudinal bond and the GDP-GTP associated energy difference of previous published BD and TK models

| Study | $U_{\text{lat}}$ | $U_{\text{long}}$ | $\Delta G_{\text{lat}}^0$ | $\Delta G_{\text{long}}^0$ | $\Delta\Delta G^0$ | Probability |
| --- | --- | --- | --- | --- | --- | --- |
| VanBuren *et al.* (2002) |  |  | -3.2 to -5.7 | -6.8 to -9.4 | 2.1-2.5 | < 0.02 |
| VanBuren *et al.* (2005) |  |  | -3.2 to -5.7 | -9.4 to -6.8 | 3.7 to 4.2 | < 0.02 |
| Gardner *et al.* (2011) |  |  | -4.5 to -5 | -9.5 | - | - |
| Margolin *et al.* (2012) |  |  | -0.4 to 1.4 | -11 to -18 | 1.75* | 0.18 |
| Coombes *et al.* (2013) |  |  | -5.7 | -7.2 | 3.3 | < 0.02 |
| Castle *et al.* (2013) | -6.8 | -20.4 | - | - | - | - |
| Zakharov *et al.* (2015) | -9.1 | -15.5 | - | - | 5.5 | < 0.02 |
| Piedra *et al.* (2016) | - | - | - | -5.8 | 3.9** | < 0.02 |
| Castle *et al.* (2017) | - | - | - | - | 3.6 | < 0.02 |
| McIntosh *et al.* (2018) | -5.3 | -16.6 | - | - | 1 | 0.31 |

All energy values are in $k_B T$
\* The maximum value of $\Delta\Delta G^0$ found within different cases of lateral neighbor in this study
\*\* This value is obtained by converting a 50-fold weakening effect in the kinetic rates to energy

### Brownian dynamics simulations of the lateral bond potential obtained via molecular dynamics imply a weak lateral bond

MD simulations’ interaction energy profiles were used to run BD simulations of two dimers in solution (Figure 6). Upon binding of a tubulin subunit, the dimer loses rotational and translational entropy as well as atomic-level entropy of the flexible side chains and loops in the protein. MD simulations can only sample the atomic-level entropy reduction due to time limitation in each umbrella window sampling (< 30 ns) whereas BD simulations can calculate the rigid body entropy cost of the coarse-grained subunits due to longer simulation times (~1ms). Thus, to fully relate the energy outputs of MD simulations to BD simulations, we calculated and subtracted rigid body entropy as a function of the reaction coordinate in MD simulations, so that the PMF would be closer to the input potential of interaction of the BD (U_BD_) with zero rigid body entropy. For rigid body entropy correction of the PMFs, Shannon entropy was calculated for each MD simulation as a function of rotational and translational degrees of freedom (Methods, Fig. S8 in the Supporting Material).

**Figure 6.**
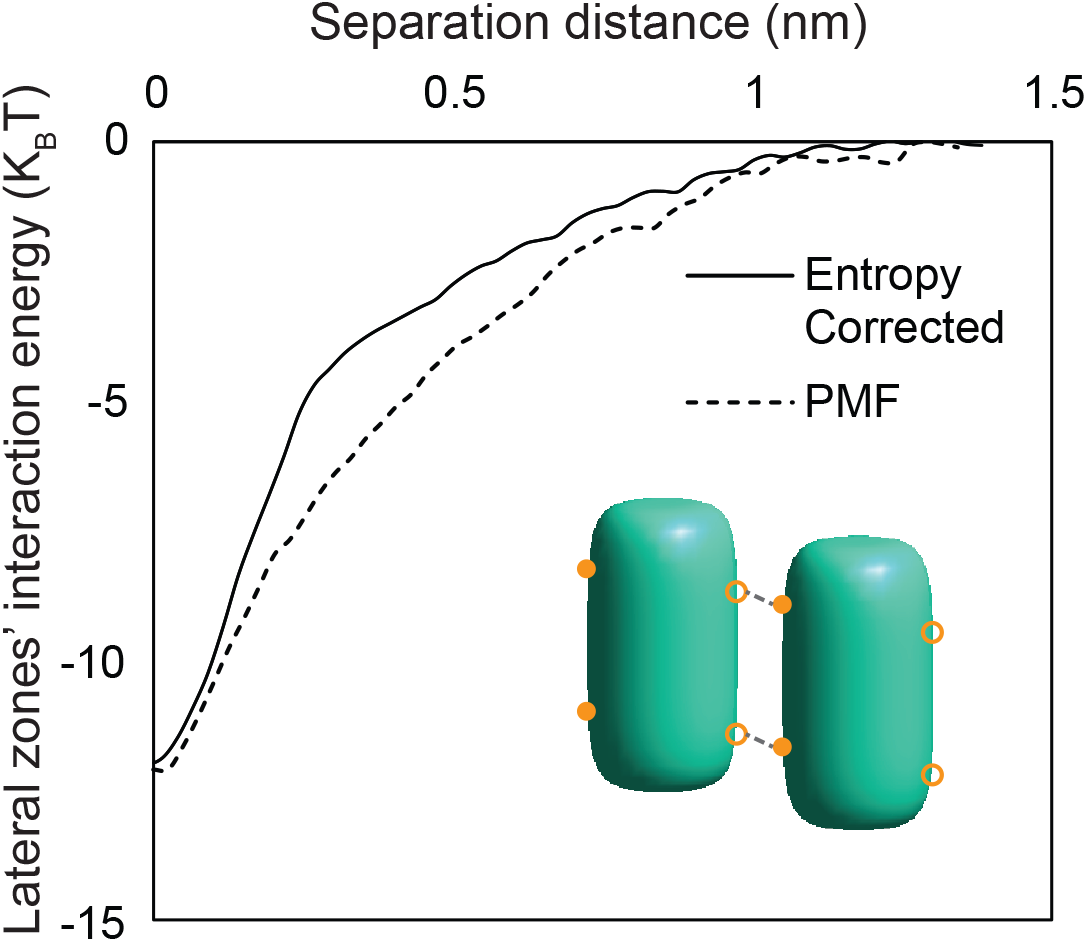
Interaction potential (U_BD_) input for GDP-tubulin from MD simulations used as lateral zone interaction energies in BD simulations. Dashed line shows the Shannon entropy-corrected energy profile and solid line shows the initial PMF resulted from MD simulation. Distance is measured from surface to surface of the rigid body dimers in BD simulations.

The BD simulations used here were modified from a previously developed model (38) to simulate free tubulins instead of a microtubule lattice. Lateral association was only allowed for the dimers through two lateral zones, similar to the previous model. Instead of using a simple harmonic potential for the lateral interaction, the estimated lateral energy landscape from MD simulations (U_MD_=PMF + TS_rigid_) was used. Since MD simulations cannot sample the rigid body entropy sufficiently due to the practical computing limits on time and length scales, we can benefit from BD simulations to obtain the full entropy cost of the lateral binding. Kinetic rates and full entropy costs were calculated for different nucleotide states (Table 4), indicating that lateral bond, by itself, is weak, unfavorable, and shortly broken after ~100ns. This is in line with the results previously obtained (38) that longitudinal association is necessary to make the lateral bond stable. Hence, the combination of the lateral bond along with the longitudinal bond makes assembly dynamics feasible in a manner previously described in thermo-kinetic (TK) terms that predict cellular level (µm scale) dynamics at time scales of minutes (35, 46). The TK modeling outputs, such as MT growth rate, shortening rate, catastrophe frequency, and rescue frequency, can be then used as inputs to cell-level models to predict dynamics and microtubule distributions associated with cell functions (47, 69–74), which allows us to now connect MD to BD to TK to CL models (Fig. 1).

**Table 4.**
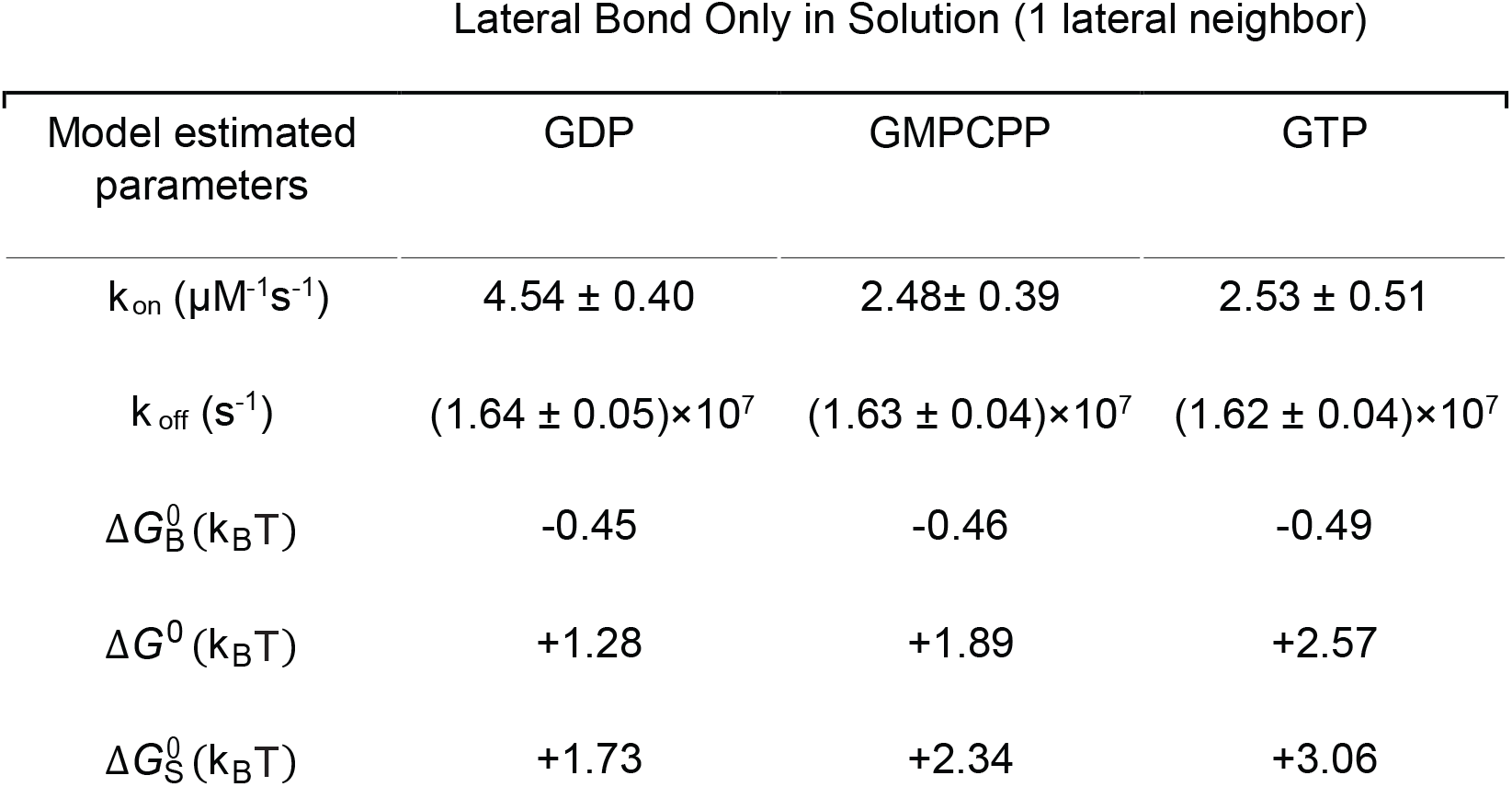
BD model outputs for a dimer in solution with one lateral neighbor with three different potential inputs

## Conclusions

A key challenge in biology is linking atomistic level information obtained by structural analysis to cellular level dynamics and function. In this study, we established a final link in an integrated multi-scale modeling chain that intertwines MD-BD-TK-CL modeling approaches for studying microtubules that connects information from atoms in crystal structures to microtubule dynamics in cells. Thus, we can seamlessly connect from Å length and fs time scales to micrometers length and minutes time scales, a span of 5 orders of magnitude in the spatial domain and 17 orders of magnitude in the temporal domain (Fig. 1). We used this approach to address a key question about the role of the lateral bond in MT stability. We found that tubulin dimers interact with each other laterally with a non-negligible force (~30-100 pN) at an edge-to-edge distance of 0.5 nm with an average strength of ~11 k_B_T, with ex estimated stiffness of ~0.2-0.5 pN/nm, regardless of nucleotide state. BD simulations revealed that the lateral bond is too weak, with a high rate of dissociation, ~10^7^ s^−1^, to be a significant pathway to MT assembly by itself. In fact, the BD simulations suggest that, given the full entropy cost of lateral binding, the total standard Gibbs free energy change of binding is unfavorable for the lateral bond by itself. This is consistent with the hypothesis that lateral bond formation is accompanied by ordering of the flexible lateral loops such as M-loop and is entropically unfavorable despite a number of favorable non-covalent interactions (40). Since dynamic instability is a key feature of microtubule assembly and vital to microtubule function, we investigated whether the lateral bond strength was nucleotide-dependent and found that it was not (<2 k_B_T). We propose that there are two possible scenarios. Either the energetic difference driving the dynamic behavior of microtubules can be incorporated into the longitudinal bond or else into the preferred bending angle in the lattice. Further MD simulations along those reaction coordinates can help us discriminate between these two possibilities.

The MD simulation findings in this study are supported by previous MD simulations of tubulin, showing that the structure of tubulin is significantly stable (simulated up to ~5μs) and the straight tubulin structure with zero lateral neighbors has a tendency for modest intradimer bending, independent of its nucleotide state (31, 33, 63, 75). The energy profiles from MD simulations in this paper are remarkably well aligned with our previous estimates based on integrated BD and TK modeling constrained by experimental kinetic rates and MT tip structure observed from assembly dynamics *in vitro* and *in vivo* (35, 37, 67). Given that the previous work started with the TK modeling at the molecular-organelle level and from there led to BD modeling, the consistency across studies increases confidence in the present estimates and further supports our conclusion that there is sufficient sampling in our MD simulations. The consistency between studies allays two concerns about the MD calculations. First, our simulation results used as a starting point a tubulin structure with kinesin bound, which may have some modest effects on the initial structure of tubulin (76). We note, however, that the equilibration MD runs reached steady-state in ~50 ns. Second, the umbrella sampling is performed on a specific chosen reaction coordinate, for which there is some uncertainty. We note, however, that the BD sims of tubulins unbinding largely follow the reaction coordinate used in the PMF calculation.

Additionally, with further MD calculations to incorporate the longitudinal bond with the lateral bond, the resulting PMFs can be combined to perform BD simulations whose outputs are k_on_ and k_off_ for single tubulin addition and loss in a thermo-kinetic model of microtubule assembly. The TK model can then be used to investigate microtubule tip structure and assembly rates in growth and shortening phases. For example, assembly rates can be used to investigate the cell-level distribution of MTs in cells such as neurons (47), consisting of numerous MTs stochastically undergoing dynamic instability. Hence, TK modeling combined with cell-level modeling can establish a direct connection with *in vivo* experimental results and large-scale cellular processes. In conclusion, our multi-scale approach provides a systematic model in which a small change in tubulin structure, due to mutation, post-translational modification, binding of a microtubule-targeting drug or microtubule associated proteins, can be connected to microtubule dynamics. Thus, the approach outlined here creates a seamless spatiotemporal connection between MD, BD, TK, and CL modeling to scale from Å to μm’s and fs to minutes, to address longstanding questions regarding the origin of MT dynamic instability.

## Author Contributions

Conceptualization, M.H. and D.J.O.; Methodology, M.H., J.S. and D.J.O.; Software, M.H. and B.T.C; Analysis, M.H., and D.J.O.; Writing – Original Draft, M.H. and D.J.O.; Writing – Review & Editing, all authors; Visualization, M.H.; Supervision, J.S, and D.J.O.; and Funding Acquisition, J.S, and D.J.O.

## Acknowledgements

The authors thank Dr. Alan M. Grossfield and Dr. David Sept for their advice and helpful discussions in preparing the paper. This study was supported by National Institutes of Health under award number R01-AG053951 and the Institute for Engineering in Medicine (IEM) award at the University of Minnesota to DJO. The authors acknowledge the Extreme Science and Engineering Discovery Environment (XSEDE) Comet at the San Diego Supercomputing Center (SDSC) through allocation MCB160060 and the Minnesota Supercomputing Institute (MSI) at the University of Minnesota for providing resources that contributed to the research results reported within this paper.

## Supporting Citations

References (75-83) appear in the Supporting Material.

