## Supporting Material for "Multi-scale Computational Modeling of Tubulin-Tubulin Lateral Interaction"

### Supporting Methods

#### Optimization of umbrella sampling parameters for enhanced sampling

Choosing the right reaction coordinate is one of the main challenges of umbrella sampling since one must manually input the proper reaction coordinate to be sampled during the simulations. For this purpose, we simulated via Brownian dynamics (BD) the lateral binding of two tubulin dimers and estimated the binding efficiency as a function of the dimer orientations with respect to each other. Doing so gave us insight into the most probable binding reaction coordinate, including the three relative rotation angles of the dimers when they bind to each other. Fig. S1 shows the binding efficiency of tubulin dimers as a function of three angles of rotation relative to each other when they are 1.5 nm center of mass (COM-to-COM) distance apart, which is within the range of the potential of mean force. The results indicates that zero rotation angle has the highest binding efficiency, meaning that if the dimers bind successfully, they are most probable to be perfectly aligned with each other. Therefore, choosing a COM-to-COM distance as a reaction coordinate for umbrella sampling is reasonable and not far from their binding path as estimated by BD.

We performed an analysis of the effect of the biasing potential stiffness on the distribution of the reaction coordinate to choose the best stiffness, i.e. one that is not so stiff that it limits the fluctuations to a narrow window without any overlap between windows but also one that is not so soft that there will be a large correlation between the atoms' movements in each window. Fig. S2 demonstrates the effects on window overlap with various stiffnesses. The $K_{bias}$=10 kcalmol$^{-1}$Å$^{-2}$ yields the best sampling results and so was chosen for all umbrella sampling simulations.

#### Potential of mean force convergence

To ensure the sampling convergence of the umbrellas sampling simulations, we increased the sampling time of each window incrementally and checked for convergence of the PMF, as shown in Fig. S3 and Table S1 and also, repeated the simulations with a different initial condition to obtain ten replicates total. Having multiple replicates samples the variability of energy distribution to a higher degree. Since all of the windows have good convergence (<1.5 $k_BT$ error determined by bootstrap error of the PMFs) after 30ns of sampling, we did not continue sampling them further. We also examined the effect of increased ion (KCl) concentration on PMF change in Fig. S4.

#### Hydrodynamic effects in MD simulations and BD simulations

Long-range hydrodynamic effects of particles binding in a solvent can lead to effects on diffusive motion of the particles thus affecting their diffusion coefficient. The classic formula of Stokes-Einstein-Sutherland (SES) for diffusion coefficient $D_0 = k_B T / 6\pi\eta a$ ($k_B T$ is the temperature, $\eta$ is solvent shear viscosity, and $a$ is the particle radius) was developed for dilute systems and does not represent the particle diffusivity accurately in concentrated dispersions (1). When particles come into contact, the viscosity of the solvent can deviate from the bulk value significantly (2, 3).

We investigated the diffusion coefficient of one tubulin dimer extracted from the COM's mean-squared-displacement in three 200ns equilibrium simulations ($D_{MD}$) and compared it to the value from the SES relationship ($D_{SE}$ ≈100 $\mu m^2 s^{-1}$) (Fig. S5). The diffusion coefficient in MD was found to be 10-fold lower than $D_{SES}$. This also reveals that despite the high dissociation rates of BD simulations (~60ns dissociation time), we do not observe any indication of dissociation of tubulin dimers in 3 replicates of 200 ns equilibrium runs. We first considered that the discrepancy could be addressed by increasing the periodic simulation box size so that the ratio of particle hydrodynamic radius to box length (R/L) decreases and long-range hydrodynamic effects of solvent declines, as suggested by previous MD studies of water diffusion (4–7). Nevertheless, increasing the box size did not have a significant effect within our timescale (~200ns). We suspect that our short time scale is the root of this deviation from SE law and a longer time scale is required for this relatively large system to reach the continuum diffusion limit. Short-time diffusion is highly dependent on hydrodynamic flows surrounding the particle and is an accumulation of local small-displacement mobility of the particle. This "Short-time" scale is defined relative to the structural relaxation time $\approx R^2/D_{SES}$ , which is ~0.15 µs for the tubulin structure. Therefore, it is not expected that the generalized SES relation would hold for such a time-scale of 200ns (1, 8, 9). Thus, the lower diffusion of the tubulin dimer observed in MD due to limited time-scale confirms the need for a multi-scale approach where larger scale simulations complement the MD results to obtain kinetic information accurately. In contrast to kinetics, we can rely on thermodynamics properties being extracted from MD simulations such as potential of mean force since we can ensure their convergence through time via enhanced sampling methods such as umbrella sampling.

In our BD simulations, we used a constant value for the diffusion coefficient of the dimers, derived from SES formula, for a simpler model. To understand how much the hydrodynamics affect tubulin's kinetic rates, we used the Honig, Roebersen, and Wiersema estimate (2) for the drag coefficient as a function of both large and small distances.

Table **S4** summarizes the hydrodynamic effects on the modeling results. Since binding is a combination of distance and alignment of the lateral zones (rotation), it is not surprising that the on-rate is not affected by hydrodynamics significantly. Higher drag coefficients at shorter distances create higher correlation in the movements of the dimers. Thus, the dimers that are in alignment at further distances are actually more probable to remain that way at shorter distances and not be affected by hydrodynamics. This is also confirmed by the fact that the binding efficiency is not influenced by hydrodynamics. Off-rate, on the other hand, is defined as the time spent within the distance criterion for separation. Hence, it is expected to be significantly reduced (~50%) by a lower diffusion coefficient as a result of the hydrodynamics. However, due to the lateral bond being weak, and therefore having a high off-rate, hydrodynamic effects on the off-rate are amplified relatively. In another run of BD simulation with both longitudinal and lateral bond present (data not shown), the hydrodynamic effects on the off-rate is weakened as well (~35%) due to stabilization by the presence of longitudinal bond.

### Supporting Movie Descriptions

**Movie S1. MD simulation of equilibration of laterally-paired GDP-tubulins with water molecules and ions (200ns) and key interacting lateral residues.** Protein structure is aligned with respect to the initial crystal structure to eliminate rigid body displacements and rotations. Water, nucleotides and ions are not shown for better visualization. Outside view of the microtubule is shown. The two key lateral interactions are highlighted (right), 1) H3 helix: H9-S10 loop (light purple), and 2) M-loop: H1'-S2 loop and H2-S3 loop (dark purple).

**Movie S2. MD simulation of equilibration of laterally-paired GMPCPP-tubulins with water molecules and ions (200ns) and key interacting lateral residues.** Protein structure is aligned with respect to the crystal structure to eliminate rigid body displacements and rotations. Water, nucleotides and ions are not shown for better visualization. Outside view of the microtubule is shown. The two key lateral interactions are highlighted (right), 1) H3 helix: H9-S10 loop (light purple), and 2) M-loop: H1'-S2 loop and H2-S3 loop (dark purple).

**Movie S3. MD simulation of equilibration of laterally-paired GTP-tubulins with water molecules and ions (200ns) and key interacting lateral residues.** Protein structure is aligned with respect to the crystal structure to eliminate rigid body displacements and rotations. Water, nucleotides and ions are not shown for better visualization. Outside view of the microtubule is shown. The two key lateral interactions are highlighted (right), 1) H3 helix: H9-S10 loop (light purple), and 2) M-loop: H1'-S2 loop and H2-S3 loop (dark purple).

### Supporting Figures

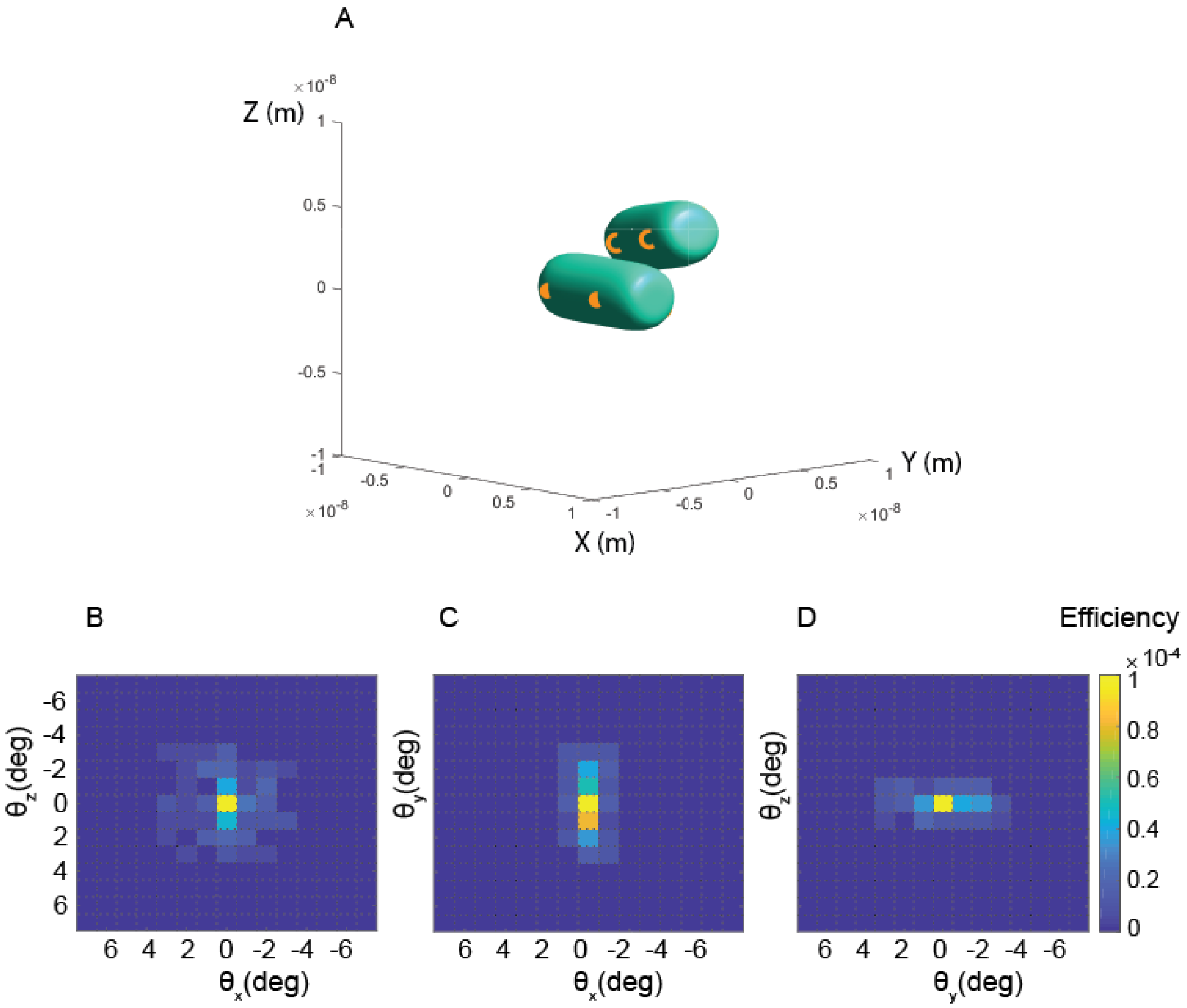


**Figure S1.** Efficiency of lateral binding for different orientations of tubulin dimers in Brownian dynamics simulation. One dimer's position and rotation is fixed at the beginning of the simulation and the other dimer is randomly positioned around it within 1.5 nm of center-to-center distance. (A) Illustration of relative position and orientation of the dimers as an example, (B-D) Binding efficiency maps for different relative angles of rotation of dimers.

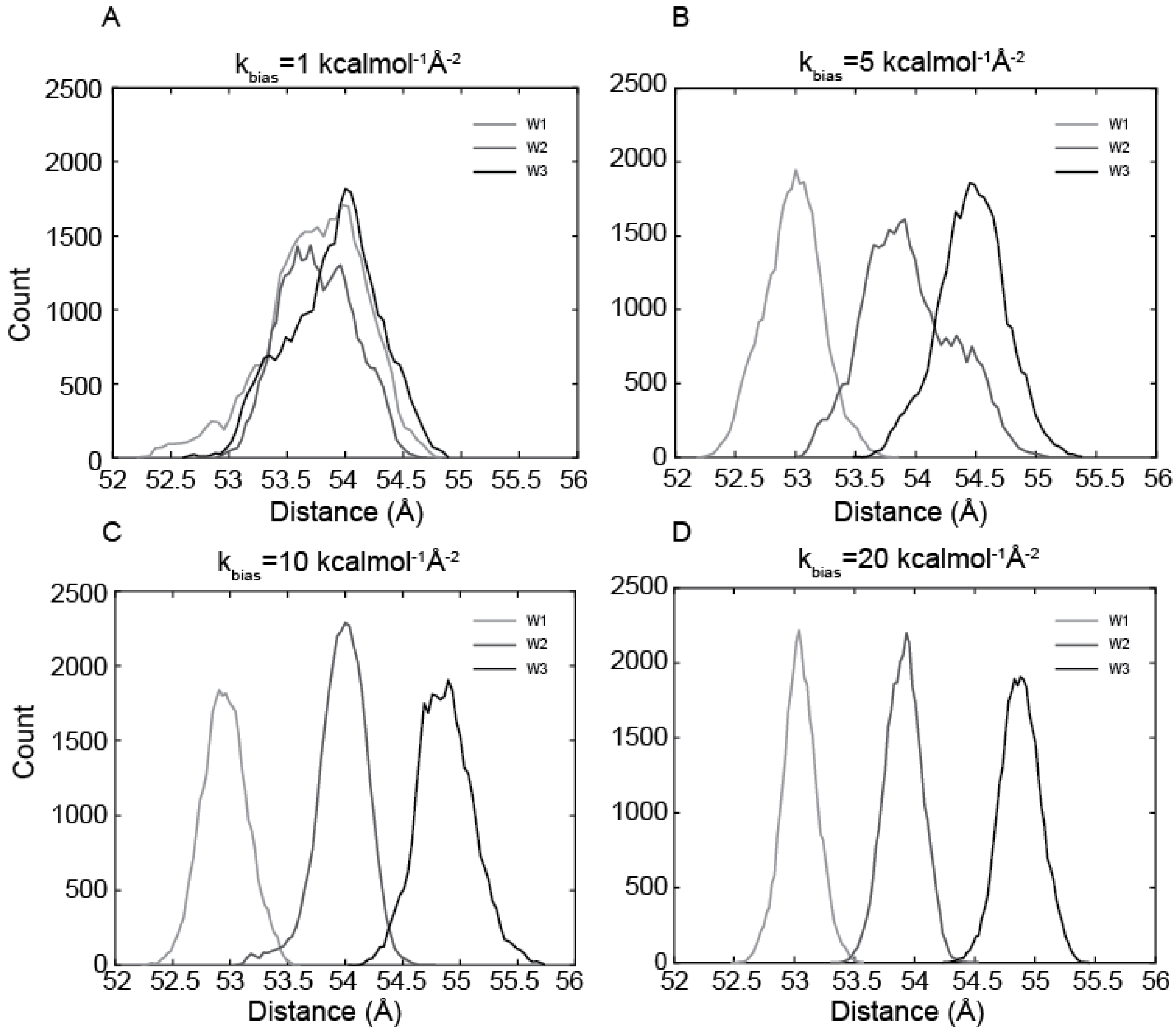


**Figure S2.** Effect of biasing potential stiffness ($k_{bias}$) on the overlap of histograms of adjacent umbrella windows near the minimum potential. (A) $k_{bias}$=1 kcal mol$^{-1}$Å$^{-2}$, (B) $k_{bias}$=5 kcal mol$^{-1}$Å$^{-2}$, (C) $k_{bias}$=10 kcal mol$^{-1}$Å$^{-2}$, and (D) $k_{bias}$=20 kcal mol$^{-1}$Å$^{-2}$.

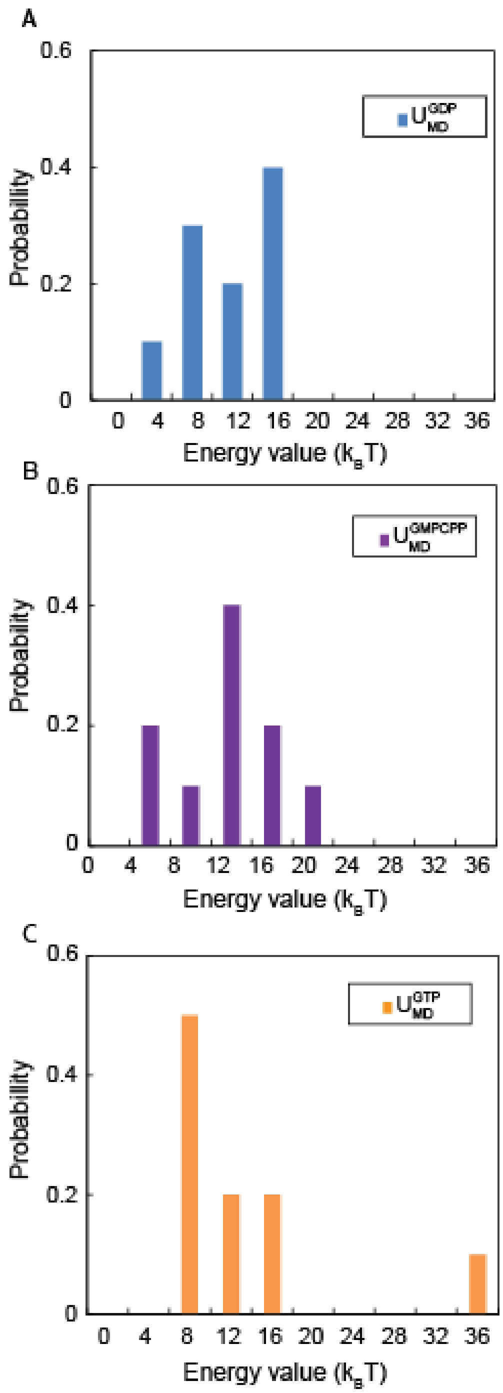


**Figure S3.** The distribution of PMF well-depth ($U_{MD}$) values obtained from different replicates of MD simulations are shown for each nucleotide states for (A) $U_{MD}^{GDP}$ , (B) $U_{MD}^{GDP}$ , and (C) $U_{MD}^{GTP}$ .

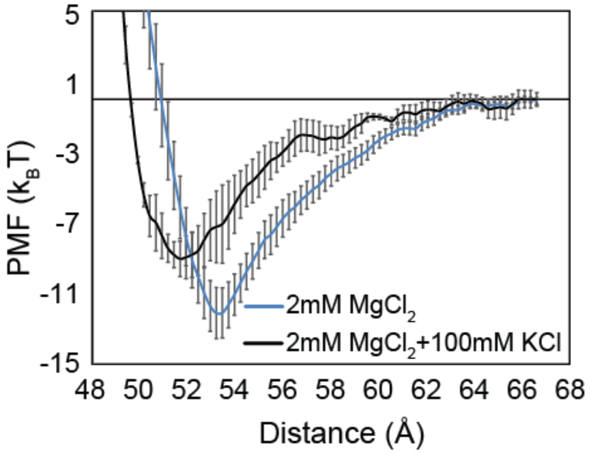


**Figure S4.** Salt concentration effects on PMF of tubulin lateral interaction. (A) Average PMF of 10 and 3 replicates for 2mM $MgCl_2$ and 2mM $MgCl_2$+100mM KCl cases, respectively. Error bars are ± standard error of the mean. (B) Well-depth, potential minimum, binding and half-force radii are calculated for different salt concentrations. P-values are calculated by Kruskal-Wallis test.

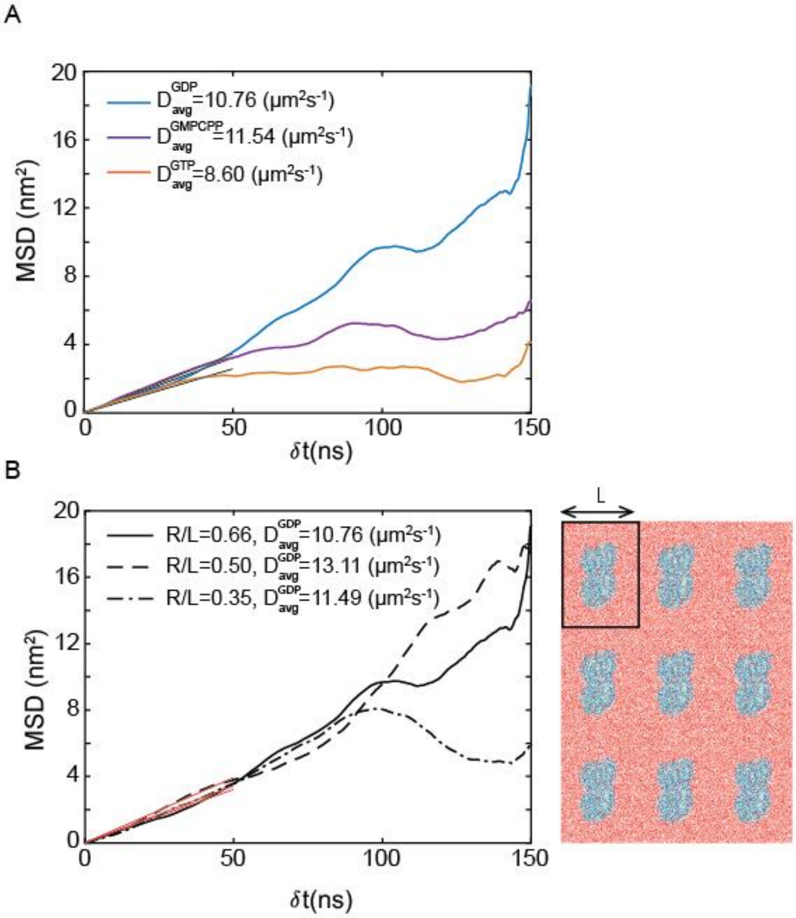


**Figure S5**. Deviation of the diffusion coefficient of tubulin in solution calculated in MD simulations from the Stokes-Einstein-Sutherland limit. (A) Mean squared displacement of a tubulin dimer in solution for different nucleotide states and resulting diffusion coefficient for a 150ns production run. (B) Diffusion coefficient of tubulin dimer in water measured for three different periodic box sizes. R indicates the calculated hydrodynamic radius of tubulin and L is the average simulation box length as shown in the right.

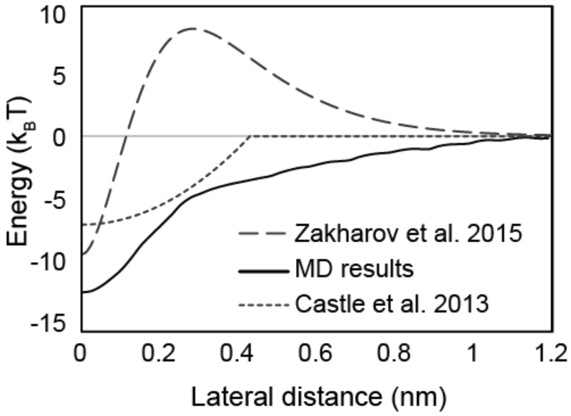


**Figure S6.** Comparison of MD lateral interaction energy profile to previously published energy profiles from Brownian dynamics models (10, 11).

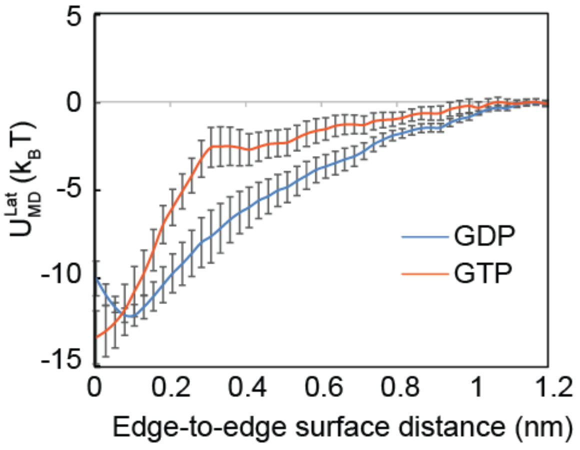


**Figure S7.** Minimum shift of the lateral potential incorporated in the input lateral potentials for BD simulations of two tubulin dimers in solution for GDP- and GTP-tubulin.

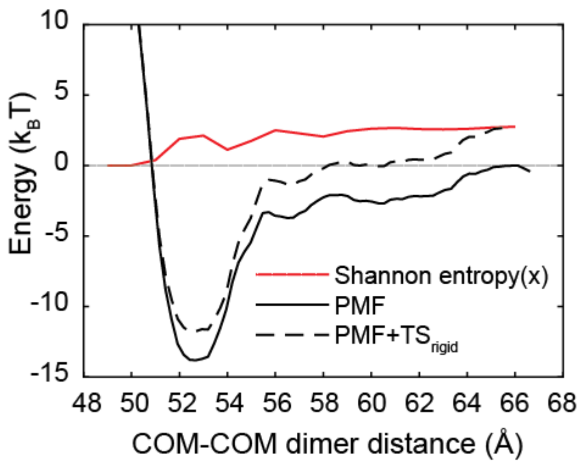


**Figure S8.** Shannon entropy correction of the PMFs for rigid body motion of tubulin dimers indicates a slight change in the potential of mean force values. Shannon entropy as a function of x is shown in red, x being the dimer distance. Original PMF is shown in black ($U_{MD}$) and entropy-corrected PMF ($U_{BD}$) is shown as a dashed black line.

### Supporting Tables

**Table S1.** The convergence of the PMF values are shown as a function of window sampling time and different initial conditions in different replicates. (A) GDP-tubulin, (B) GMPCPP-tubulin, (C) GTP-tubulin. A tolerance of 1 $k_BT$ was considered for choosing the sampling time.

A

| Replicate | $\Delta G^0_{GDP}$ ($k_BT$) 15 ns sampling | 21 ns sampling | 30 ns sampling |
|---|---|---|---|
| 1 | 5.5 ± 2.9 | 6.8 ± 1.7 | 7.6 ± 1.5 |
| 2 | 9.6 ± 2.6 | 6.8 ± 2.2 | 8.3 ± 1.6 |
| 3 | 11.4 ± 2.4 | 11.7 ± 1.9 | 12.5 ± 1.7 |
| 4 | 12.6 ± 2.4 | 12.1 ± 1.7 | 12.8 ± 1.5 |
| 5 | 17.7 ± 2.8 | 18.3 ± 1.8 | 15.0 ± 1.5 |
| 6 | 11.0 ± 2.7 | 9.0 ± 2.0 | 10.8 ± 1.9 |
| 7 | 7.5 ± 2.2 | 6.9 ± 1.9 | 7.2 ± 1.4 |
| 8 | 13.5 ± 1.9 | 14.7 ± 1.9 | 13.9 ± 1.5 |
| 9 | 18.4 ± 2.2 | 17.8 ± 1.9 | 14.2 ± 1.7 |
| 10 | 10.7 ± 2.5 | 9.7 ± 1.9 | 9.6 ± 1.6 |

B

| Replicate | $\Delta G^0_{GMPCPP}$ ($k_BT$) 15 ns sampling | 21 ns sampling | 30 ns sampling |
|---|---|---|---|
| 1 | 16.1 ± 2.2 | 15.6 ± 1.7 | 14.2 ± 1.6 |
| 2 | 9.1 ± 2.3 | 7.7 ± 2.0 | 8.3 ± 1.6 |
| 3 | 19.4 ± 2.4 | 19.4 ± 1.6 | 15.9 ± 1.4 |
| 4 | 9.9 ± 2.4 | 9.2 ± 1.9 | 9.5 ± 1.6 |
| 5 | 24.8 ± 2.6 | 23.8 ± 1.8 | 20.9 ± 1.5 |
| 6 | 20.1 ± 2.5 | 19.2 ± 1.9 | 15.8 ± 1.5 |
| 7 | 4.8 ± 3.1 | 5.8 ± 2.3 | 6.1 ± 1.6 |
| 8 | 10.3 ± 2.2 | 8.6 ± 2.1 | 6.7 ± 1.7 |
| 9 | 11.7 ± 2.4 | 10.6 ± 1.9 | 9.9 ± 1.6 |
| 10 | 15.8 ± 2.4 | 14.9 ± 1.5 | 12.9 ± 1.4 |

C

| Replicate | $\Delta G^0_{GTP}$ ($k_BT$) 15 ns sampling | 21 ns sampling | 30 ns sampling |
|---|---|---|---|
| 1 | 25.2 ± 2.2 | 23.8 ± 1.9 | 20.7 ± 1.5 |
| 2 | 32.1 ± 2.7 | 32.0 ± 2.1 | 30.8 ± 1.9 |
| 3 | 12.4 ± 3.0 | 12.2 ± 2.0 | 11.6 ± 1.6 |
| 4 | 16.8 ± 2.4 | 12.9 ± 2.0 | 13.2 ± 1.5 |
| 5 | 7.7 ± 3.0 | 9.1 ± 2.1 | 9.1 ± 1.7 |
| 6 | 18.2 ± 2.4 | 15.5 ± 2.1 | 12.4 ± 1.6 |
| 7 | 11.8 ± 2.7 | 8.8 ± 1.7 | 8.5 ± 1.3 |
| 8 | 15.2 ± 2.3 | 14.4 ± 1.9 | 13.6 ± 1.8 |
| 9 | 14.7 ± 2.8 | 12.4 ± 1.8 | 10.0 ± 1.5 |
| 10 | 13.8 ± 1.7 | 12.2 ± 1.8 | 12.7 ± 1.5 |

**Table S2.** BD simulation results for incorporating GDP-tubulin minimum shift in lateral potential.

Lateral Bond in solution (1 lateral neighbor)

| Model estimated parameters | GTP-tubulin | GDP-tubulin With minimum shift |
|---|---|---|
| $k_{on}$ ($\mu M^{-1}s^{-1}$) | 2.53 ± 0.51 | 5.25± 0.89 |
| $k_{off}$ ($s^{-1}$) | $1.62\times10^{7}$ ± 0.04 | $1.63\times10^{6}$± 0.06 |
| $\Delta G_B^0$ ($k_BT$) | -0.49 | -0.29 |
| $\Delta G^0$ ($k_BT$) | +2.57 | +1.13 |
| $\Delta G_S^0$ ($k_BT$) | +3.06 | +1.43 |

**Table S3.** Lateral PMF comparison in different ion concentrations.

| Salt concentration | Well depth ($k_BT$) ± SEM | Binding radius (nm) ± SEM | Half-force radius (nm) ± SEM | Potential minimum (nm) ± SEM |
|---|---|---|---|---|
| 2mM $MgCl_2$ | 10.95 ± 1.33 | 0.90 ± 0.06 | 0.42 ± 0.03 | 5.39 ± 0.02 |
| 2mM $MgCl_2$ +100mM KCl | 8.40 ± 1.17 | 0.82 ± 0.06 | 0.42 ± 0.02 | 5.19 ± 0.05* |

* P-values < 0.02

**Table S4.** Hydrodynamic effects on the kinetic rates in BD simulation of tubulin-tubulin lateral association.

Lateral Bond in solution (1 lateral neighbor)-GDP Tubulin

| Model estimated parameters | $D_{SE}$ | Hydrodynamic D |
|---|---|---|
| $k_{on}$ ($\mu M^{-1}s^{-1}$) | 4.54 ± 0.40 | 4.31± 0.22 |
| $k_{off}$ ($s^{-1}$) | $1.64\times10^{7}$ ± 0.05 | $8.48\times10^{6}$± 0.03 |
| $\Delta G_B^0$ ($k_BT$) | -0.45 | -1.74 |
| $\Delta G^0$ ($k_BT$) | +1.28 | +0.68 |
| $\Delta G_S^0$ ($k_BT$) | +1.73 | +2.42 |
